# Gene duplications trace mitochondria to the onset of eukaryote complexity

**DOI:** 10.1101/781211

**Authors:** Fernando D. K. Tria, Julia Brückner, Josip Skejo, Joana C. Xavier, Verena Zimorski, Sven B. Gould, Sriram G. Garg, William F. Martin

## Abstract

The last eukaryote common ancestor (LECA) lived 1.6 billion years ago^1,2^. It possessed nuclei, sex, an endomembrane system, mitochondria, and all key traits that make eukaryotic cells more complex than their prokaryotic ancestors^2–6^. The closest known relatives of the host lineage that acquired the mitochondrion are, however, small obligately symbiotic archaea that lack any semblance of eukaryotic cell complexity^7^. Although the steep evolutionary grade separating prokaryotes from eukaryotes increasingly implicates mitochondrial symbiosis at eukaryote origin^4,7^, the timing and evolutionary significance of mitochondrial origin remains debated. Gradualist theories contend that eukaryotes arose from archaea by slow accumulation of eukaryotic traits^8–10^ with mitochondria arriving late^11^, while symbiotic theories have it that mitochondria initiated the onset of eukaryote complexity in a non-nucleated archaeal host^7^ by gene transfers from the organelle^4,12–14^. The evolutionary process leading to LECA should be recorded in its gene duplications. Among 163,545 duplications in 24,571 gene trees spanning 150 sequenced eukaryotic genomes we identified 713 gene duplication events that occurred in LECA. LECA’s bacterially derived genes were duplicated more frequently than archaeal derived or eukaryote specific genes, reflecting the serial copying^15,16^ of genes from the mitochondrial endosymbiont to the archaeal host’s chromosomes prior to the onset of eukaryote genome complexity. Bacterial derived genes for mitochondrial functions, lipid synthesis, biosynthesis, as well as core carbon and energy metabolism in LECA were duplicated more often than archaeal derived genes and even more often than eukaryote-specific inventions for endomembrane, cytoskeletal or cell cycle functions. Gene duplications record the sequence of events at LECA’s origin and indicate that recurrent gene transfer from a resident mitochondrial endosymbiont preceded the onset of eukaryotic cellular complexity.

## Main text

Gene duplication is the hallmark of eukaryotic genome evolution^17^. Individual gene families^18^ and whole genomes^19,20^ have undergone recurrent duplication across the eukaryotic lineage. By contrast, gene duplications in prokaryotes are rare at best^21^ and whole genome duplications of the nature found in eukaryotes are unknown. Gene duplication is a eukaryotic trait. Its origin is of interest. In order to learn more about the onset of gene duplication in eukaryote genome evolution, we investigated duplications in sequenced genomes (see **Methods**). We plotted all duplications shared by at least two eukaryotic genomes among 1,848,936 protein-coding genes from 150 sequences eukaryotes spanning six supergroups: Archaeplastida, Opisthokonta, Mycetozoa, Hacrobia, SAR and Excavata^22^. Nearly half of all eukaryotic genes (941,268) exist as duplicates in 239,012 gene families. Of those, 24,571 families (10.3%) harbor duplicate copies in at least two eukaryotic genomes (multi-copy gene families), with variable distribution across the supergroups (**Fig. 1**). Opisthokonta harbor in total 22,410 multi-copy gene families, with the largest number by far (19,530) present among animals followed by 6,495 multicopy gene families in the plant lineage (Archaeplastida). Among the 24,571 multi-copy gene families, 1,823 are present in at least one genome from all six supergroups and are potential candidates of gene duplications tracing to LECA.

**Figure 1:**
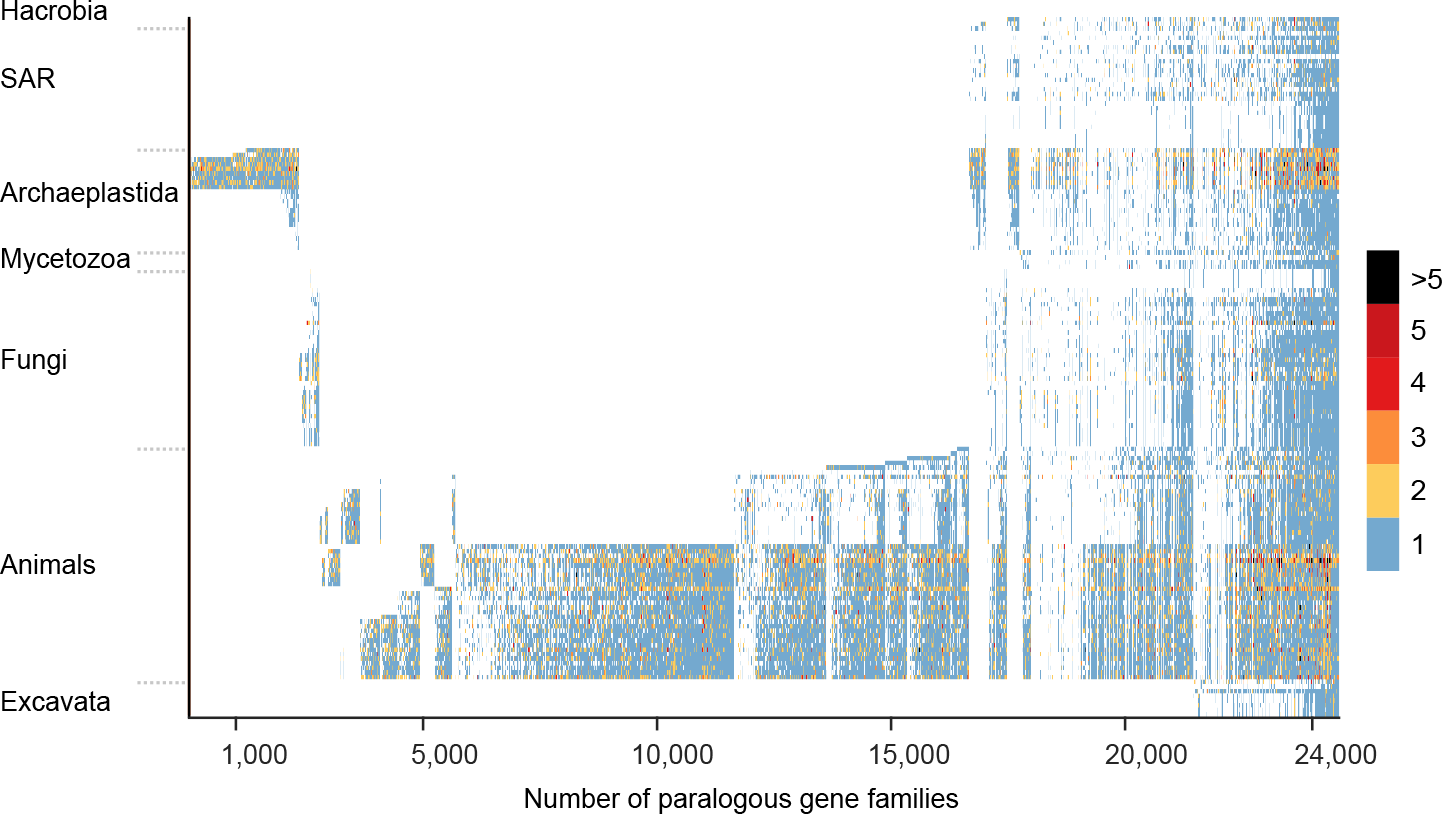
Distribution of multi-copy genes across 150 eukaryotic genomes. The protein sequences were clustered using the MCL algorithm and the resulting gene families present as multiple copies in more than one genome are plotted (see **Methods**). The figure displays the 24,571 multi-copy gene families (horizontal axis), the colored scale indicates the number of gene copies in each eukaryotic genome (vertical axis). The genomes were sorted according to a reference species tree (**Supplemental data**) and taxonomic classifications were taken from NCBI^41^. Animals and fungi together form the opisthokont supergroup.

To identify the relative phylogenetic timing of eukaryotic gene duplication events, we reconstructed maximum-likelihood trees from protein alignments for all individual multi-copy gene families. In each gene tree, we assigned gene duplications to the most recent branch possible, allowing for multiple gene duplication events if needed (see **Methods**) and permitting any branching order of supergroups. This identified 163,545 gene duplications, 160,676 of which generate paralogs within a single supergroup and an additional 2,869 gene duplication events that trace to the common ancestor of at least two supergroups (**Fig. 2a and Supplemental Table 1**). The results show that gene duplications were taking place in LECA before the eukaryotic supergroups diverged, because for 713 duplications in 475 gene families, the resulting paralogs are distributed across all six supergroups, indicated in red in **Fig. 2a**.

**Figure 2:**
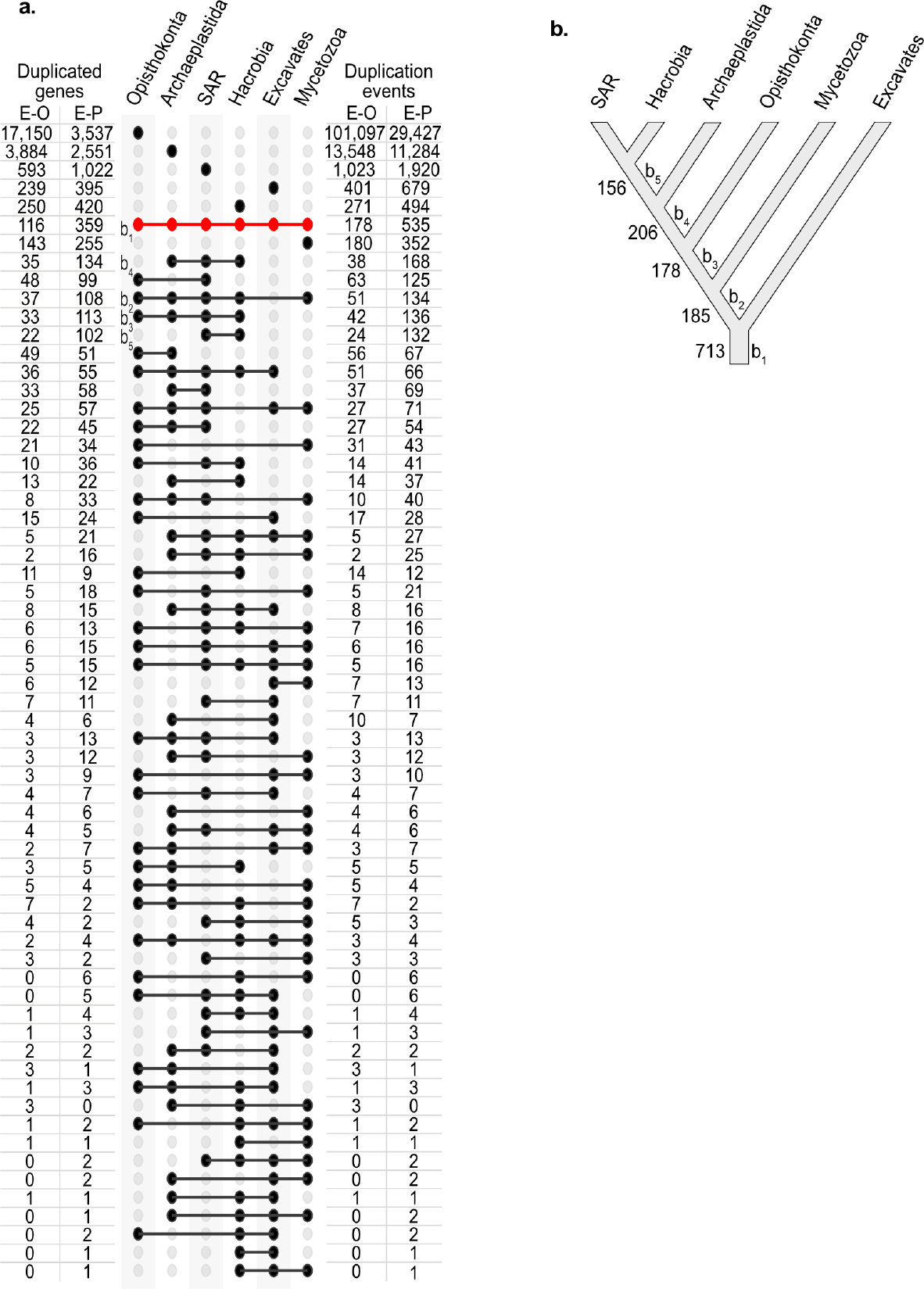
Distribution of gene duplications across six eukaryotic supergroups. **a)** The figure shows the distribution of paralogs resulting from the inferred gene duplications in eukaryotic-specific genes (E-O) and eukaryotic genes with prokaryotic homologs (E-P) (see **Methods** for details). Duplicated genes refer to the numbers of gene trees with at least one duplication assigned to the common ancestor of supergroups (filled circles in the center). Number of duplication events refers to the total number of gene duplications. Note that a gene may experience multiple gene duplications. The red row circles indicate gene trees with duplications in LECA and descendant paralogs in all six supergroups. An early study assigned 4,137 duplicated gene families to LECA but attributed all copies present in any two major eukaryotic groups to LECA^72^; in the present sample, we find 2,869 gene duplication events that trace to the common ancestor of at least two supergroups. Our stringent criterion requiring paralogue presence in all six supergroups leaves 713 duplications in 475 gene families in LECA. **b)** Rooted phylogeny of eukaryotic supergroups that maximizes compatibility with gene duplications. Duplications mapping to the five external edges are shown (b_1_, b_2_, …, b_5_). The tree represents almost exactly all possible edges containing the most duplications, the exception is the branch joining Hacrobia and SAR, which has a more supported branch uniting SAR and Opisthokonta, but the resulting subtree ((Opisthokonta,SAR),(Archaeplastida, Hacrobia)) accounts for 249 duplications, fewer than the (Opisthokonta,(Archaeplastida,(SAR, Hacrobia))) subtree shown (262 duplications). The position of the root identifies additional duplications with descendant paralogs in Excavata and other supergroup(s) that trace to LECA (**Table 1 and Supplemental Table 4**).

The six supergroups plus LECA at the root represent a seven-taxon tree in which the external edges bearing the vast majority of duplications (**Fig. 2a**). Gene duplications that map to internal branches of the rooted supergroup tree can result from duplications in LECA followed by vertical inheritance and differential loss in some supergroups, or they can map to the tree by which supergroups are related following their divergence from LECA. Branches that explain the most duplications are likely to reflect the natural phylogeny, because support for conflicting branches from random^19^ non-phylogenetic patterns are generated by independent losses. There is a strong phylogenetic signal contained within eukaryotic gene duplication data (**Fig. 2**). Among all possible internal branches, those supported by the most frequent duplications are compatible with the tree in **Fig. 2b**, which places the eukaryotic root on the branch separating Excavates^22^ from other supergroups, as implicated in previous studies of concatenated protein sequences^23,24^.

LECA’s duplications address the timing of mitochondrial origin, because different theories for eukaryote origin generate different predictions about the nature of duplications in LECA. Gradualist theories^8–11^ predict archaeal specific and eukaryote specific genes to have undergone numerous duplications during the origin of eukaryote complexity prior to the acquisition of the mitochondrion that completed the process of eukaryogenesis. In that case, bacterial derived genes would have accumulated fewer duplications in LECA than archaeal derived or eukaryote specific genes (**Fig. 3a**). Models invoking gradual lateral gene transfers (LGT) from ingested (phagocytosed) food prokaryotes prior to the origin of mitochondria^25^ also predict more duplications in archaeal derived and eukaryote specific genes to underpin the origin of phagocytotic feeding, but do not predict duplications specifically among acquired genes (whether from bacterial or archaeal food) because each ingestion contributes genes only once. By contrast, transfers from the endosymbiotic ancestors of organelles continuously generate duplications in the host’s chromosomes^15,16^, a process that continues to the present day in eukaryotic genomes^16,26^.

**Figure 3:**
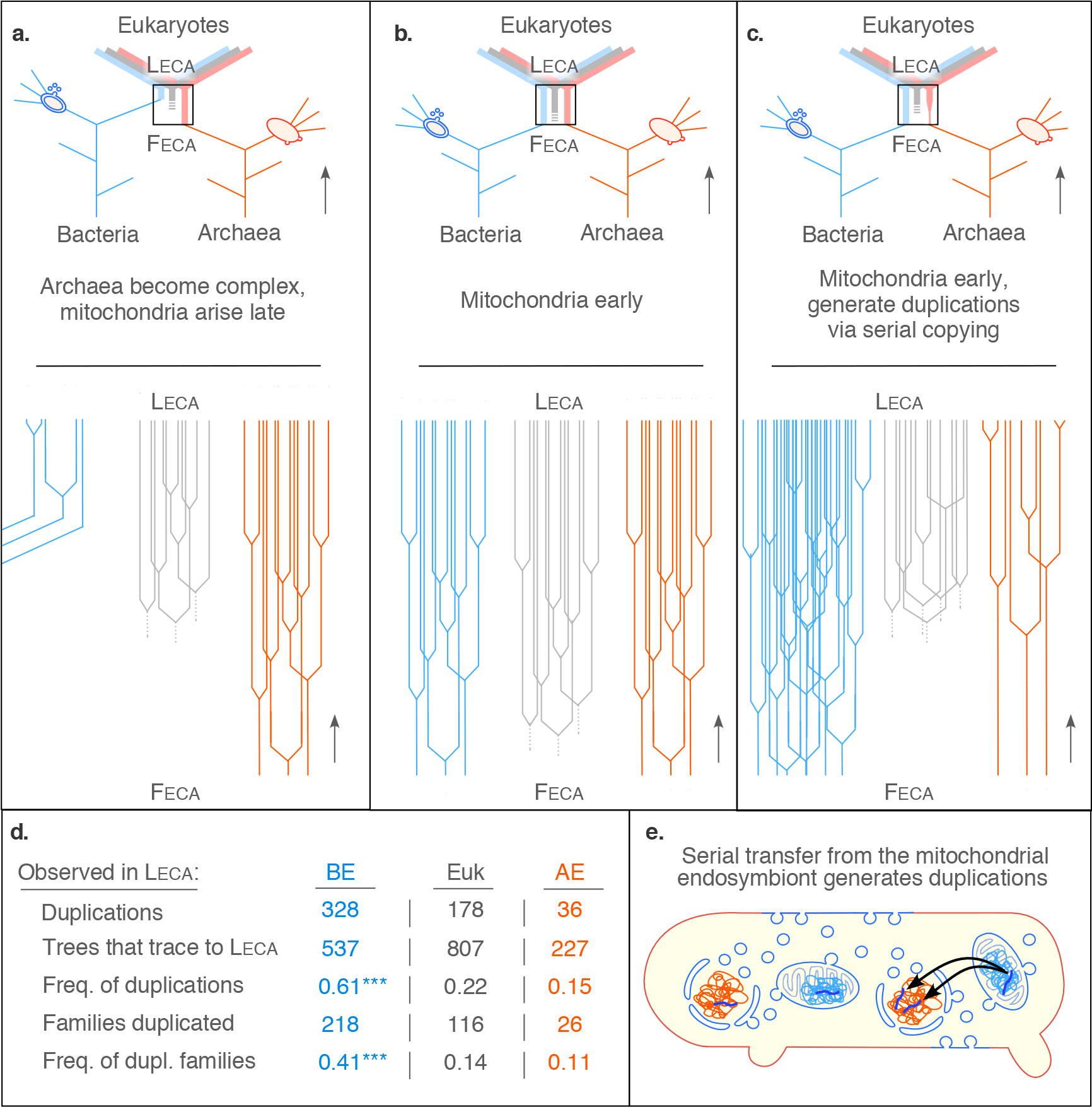
Different models for eukaryote origin generate different predictions with respect to duplications. In each panel, gene duplications during the FECA to LECA transition (boxed in upper portion) is enlarged in the lower portion of the panel. **a)** Cellular complexity and genome expansion in an archaeal host predates the origin of mitochondria. **b)** Mitochondria enter the eukaryotic lineage early, duplications in mitochondrial derived, host derived and eukaryotic specific genes occur, genome expansion affects all genes equally. **c)** Gene transfers from a resident endosymbiont generate duplications in genes of bacterial origin in an archaeal host. **d)** Observed frequencies from gene duplications that trace to LECA (see **Supplemental Table 3**). BE refers to eukaryotic genes with bacterial homologs only; AE refers to eukaryotic genes with archaeal homologs only; and Euk refers to eukaryotic genes without prokaryotic homologs. **e)** Serial gene transfers from the mitochondrion (blue components) generate duplicates in the chromosomes of the host (red components). Outer membrane vesicles of the mitochondrion^4^ and the host^7^, the former leading to lipid replacement in eukaryotes^12^ and the origin of the endomembrane system^4^, are indicated.

Symbiogenic theories posit that the host that acquired the mitochondrion was an archaeon of normal prokaryotic complexity^4,7,12–14^ and hence lacked duplications underpinning eukaryote complexity. There are examples known in which bacteria grow in intimate association with archaea^13^ and in which prokaryotes become endosymbionts within other prokaryotic cells^13^. Energetic constraints^14^ to genome expansion apply to all genes, highly expressed genes in particular, such that gene duplications in the wake of mitochondrial origin should be equally common in genes of bacterial, archaeal or eukaryote-specific origin, respectively (**Fig. 3b**). Gene transfers from resident organelles involve endosymbiont lysis and incorporation of complete organelle genomes followed by recombination and mutation^26^. In contrast to LGTs from extracellular donors, gene transfers from resident endosymbionts specifically generate duplications because new copies of the same genes are recurrently transferred^15–17^ (**Fig. 3c**). The duplications in LECA reveal a vast excess of duplications in LECA’s bacterial derived genes relative to archaeal derived and eukaryote-specific genes, respectively (**Fig. 3d**). The proportion of duplications in bacterial derived genes is fourfold and threefold higher than for archaeal derived and eukaryote specific genes.

The association of duplications tracing to LECA and genes with bacterial counterparts is significant among eukaryotic genes distributed in all six supergroups, as judged by the two-tailed Fisher’s test (p-value < 0.001, **Supplemental Table 2**). Based on the functions of duplicates (**Table 1**), the resident endosymbiont in LECA was the mitochondrion (**Fig. 3e**). Gene duplications in 48 genes with mitochondrial functions include pyruvate dehydrogenase complex, enzymes of the citric acid cycle, components involved in electron transport, a presequence cleavage protease, the ATP-ADP carrier, and 7 members of the eukaryote-specific mitochondrial carrier family that facilitates metabolite exchange between the mitochondrion and the cytosol (**Table 1**; **Supplemental Tables 3 and 4**). This indicates that canonical energy metabolic functions of mitochondria had been established in LECA, underscored by additional functions performed by mitochondria in diverse eukaryotic lineages: 11 genes for enzymes of the lipid biosynthetic pathway (typically mitochondrial in eukaryotes^4^), the entire glycolytic pathway (mitochondrial among marine algae^27^), and 10 genes involved in redox balance are found among bacterial duplicates. The largest category of duplications with annotated functions concerns metabolism and biosynthesis (**Table 1**).

**Table 1.**
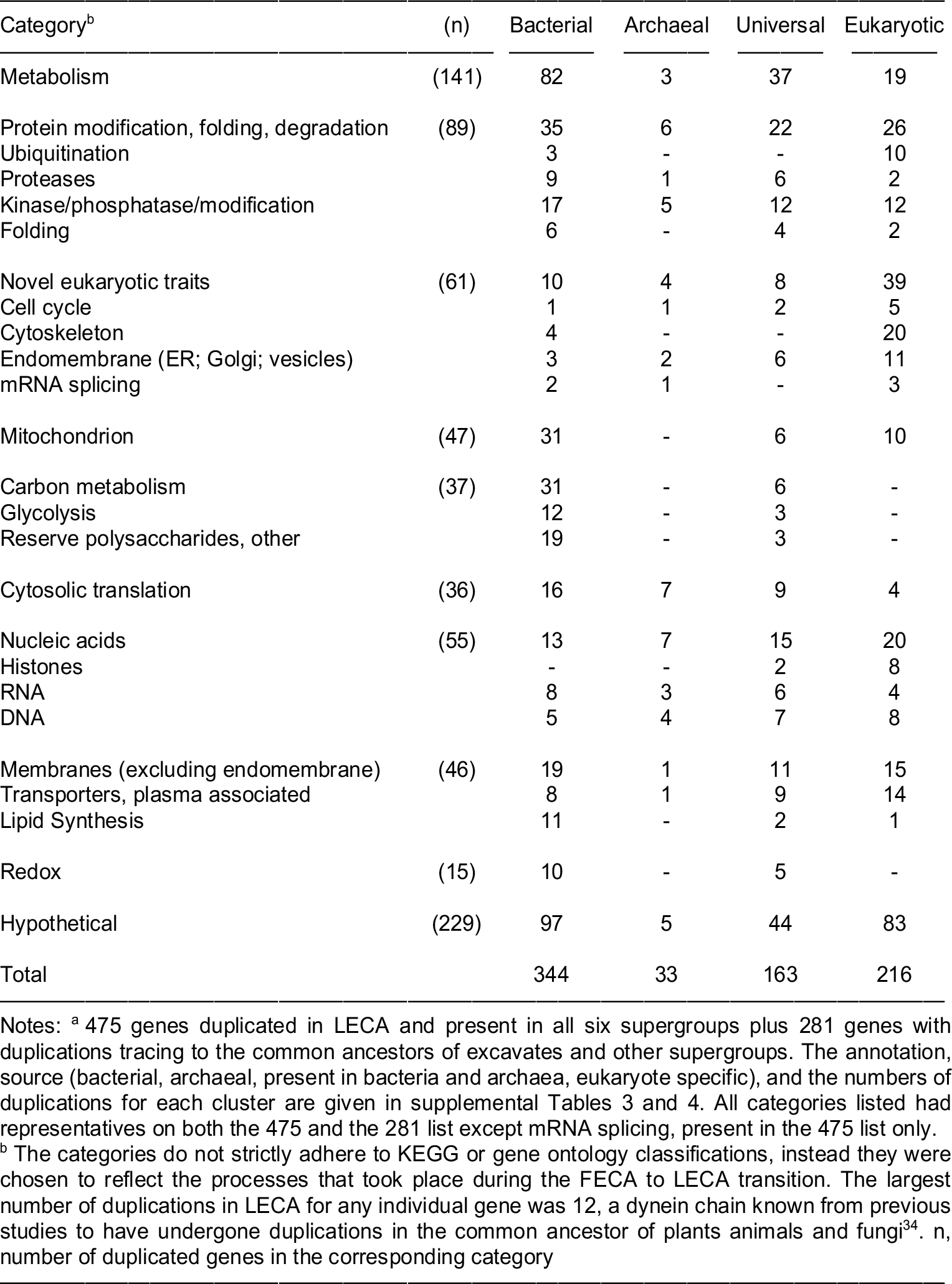
Functional categories of genes duplicated in LECA^a^.

| Category<sup>b</sup> | (n) | Bacterial | Archaeal | Universal | Eukaryotic |
| --- | --- | --- | --- | --- | --- |
| Metabolism | (141) | 82 | 3 | 37 | 19 |
| Protein modification, folding, degradation | (89) | 35 | 6 | 22 | 26 |
| Ubiquitination |  | 3 | - | - | 10 |
| Proteases |  | 9 | 1 | 6 | 2 |
| Kinase/phosphatase/modification |  | 17 | 5 | 12 | 12 |
| Folding |  | 6 | - | 4 | 2 |
| Novel eukaryotic traits | (61) | 10 | 4 | 8 | 39 |
| Cell cycle |  | 1 | 1 | 2 | 5 |
| Cytoskeleton |  | 4 | - | - | 20 |
| Endomembrane (ER; Golgi; vesicles) |  | 3 | 2 | 6 | 11 |
| mRNA splicing |  | 2 | 1 | - | 3 |
| Mitochondrion | (47) | 31 | - | 6 | 10 |
| Carbon metabolism | (37) | 31 | - | 6 | - |
| Glycolysis |  | 12 | - | 3 | - |
| Reserve polysaccharides, other |  | 19 | - | 3 | - |
| Cytosolic translation | (36) | 16 | 7 | 9 | 4 |
| Nucleic acids | (55) | 13 | 7 | 15 | 20 |
| Histones |  | - | - | 2 | 8 |
| RNA |  | 8 | 3 | 6 | 4 |
| DNA |  | 5 | 4 | 7 | 8 |
| Membranes (excluding endomembrane) | (46) | 19 | 1 | 11 | 15 |
| Transporters, plasma associated |  | 8 | 1 | 9 | 14 |
| Lipid Synthesis |  | 11 | - | 2 | 1 |
| Redox | (15) | 10 | - | 5 | - |
| Hypothetical | (229) | 97 | 5 | 44 | 83 |
| Total |  | 344 | 33 | 163 | 216 |

Notes: <sup>a</sup> 475 genes duplicated in LECA and present in all six supergroups plus 281 genes with duplications tracing to the common ancestors of excavates and other supergroups. The annotation, source (bacterial, archaeal, present in bacteria and archaea, eukaryote specific), and the numbers of duplications for each cluster are given in supplemental Tables 3 and 4. All categories listed had representatives on both the 475 and the 281 list except mRNA splicing, present in the 475 list only.
<sup>b</sup> The categories do not strictly adhere to KEGG or gene ontology classifications, instead they were chosen to reflect the processes that took place during the FECA to LECA transition. The largest number of duplications in LECA for any individual gene was 12, a dynein chain known from previous studies to have undergone duplications in the common ancestor of plants animals and fungi<sup>34</sup>. n, number of duplicated genes in the corresponding category

Many products of bacterial derived genes operate in the eukaryotic cytosol. This is because at the outset of gene transfer from the endosymbiont, there was no mitochondrial protein import machinery^12,28^, such that the products of genes transferred from the endosymbiont were active in the compartment where the genes were co-transcriptionally translated^29^. Gene transfers in large, genome sized fragments from the endosymbiont, as they occur today^16,26^, furthermore permitted entire pathways to be transferred, because the unit of biochemical selection is the pathway and its product, not the individual enzyme^30^. In the absence of upstream and downstream intermediates and activities in a pathway, the product of a lone transferred gene is generally useless for the cell, expression of the gene becomes a burden and the transferred gene cannot be fixed^30^.

The origin of mRNA splicing, a selective force at the origin of the nucleus^31^, the origin of the endomembrane system from mitochondrion derived vesicles (MDVs) of bacterial lipids^4^, and the origin of protein import in mitochondria^28^, all present in LECA, established cell compartmentation in the first eukaryote. Notably, duplicate genes of bacterial origin are also involved in the origin of eukaryotic specific traits, including the cell cycle, the cytoskeleton, endomembrane system and mRNA splicing (**Table 1**). The bacterial duplicate contribution exceeds the archaeal contribution to these categories, which are dominated by eukaryote-specific genes. Duplications in LECA depict bacterial carbon and energy metabolism in an archaeal host supported by genes that were recurrently donated by a resident symbiont, in line with the predictions of symbiotic theories for the nature of the first eukaryote^7,12,13^, but contrasting sharply with theories involving eukaryote origin from phagocytosing archaea^8–11^.

Like the nucleus, mitochondria, and other eukaryotic traits^2–7^, the accrual of gene and genome duplications distinguish eukaryotes from prokaryotes^17–21^. Gene transfers from the mitochondrion can generate duplications of bacterial derived genes. What mechanisms promoted genome-wide gene duplication at the prokaryote-eukaryote transition? Population genetic parameters such as variation in population size^6^ apply to prokaryotes and eukaryotes equally, hence they would not affect gene duplications specifically in eukaryotes, but recombination processes^31^ in a nucleated cell could. Because LECA possessed meiotic recombination^3^, it was able to fuse nuclei (karyogamy). Karyogamy in a multinucleate LECA would promote the accumulation of duplications in all gene classes and genome expansion to its energetically permissible limits^14^ because unequal crossing between imprecisely paired homologous chromosomes following karyogamy generates duplications^17–20^. At the origin of meiotic recombination, chromosome pairing and segregation cannot have been perfect from the start; the initial state was likely error-prone, generating nuclei with aberrant gene copies, aberrant chromosomes or even aberrant chromosome numbers. In cells with a single nucleus, such variants would have been lethal; in multinucleate (syncytial or coenocytic) organisms, defective nuclei can complement each other through mRNA in the cytosol^31^. Multinucleate forms are present throughout eukaryotic lineages (**Fig. 4**), and ancestral reconstruction of nuclear organization clearly indicates that LECA itself was multinucleate (**Fig. 4**). The multinucleate state enables the accumulation of duplications in the incipient eukaryotic lineage in a mechanistically non-adaptive manner, and duplications are implicated in the evolution of complexity^17–20^, as observed in the animal lineage (**Fig. 1**).

**Figure 4:**
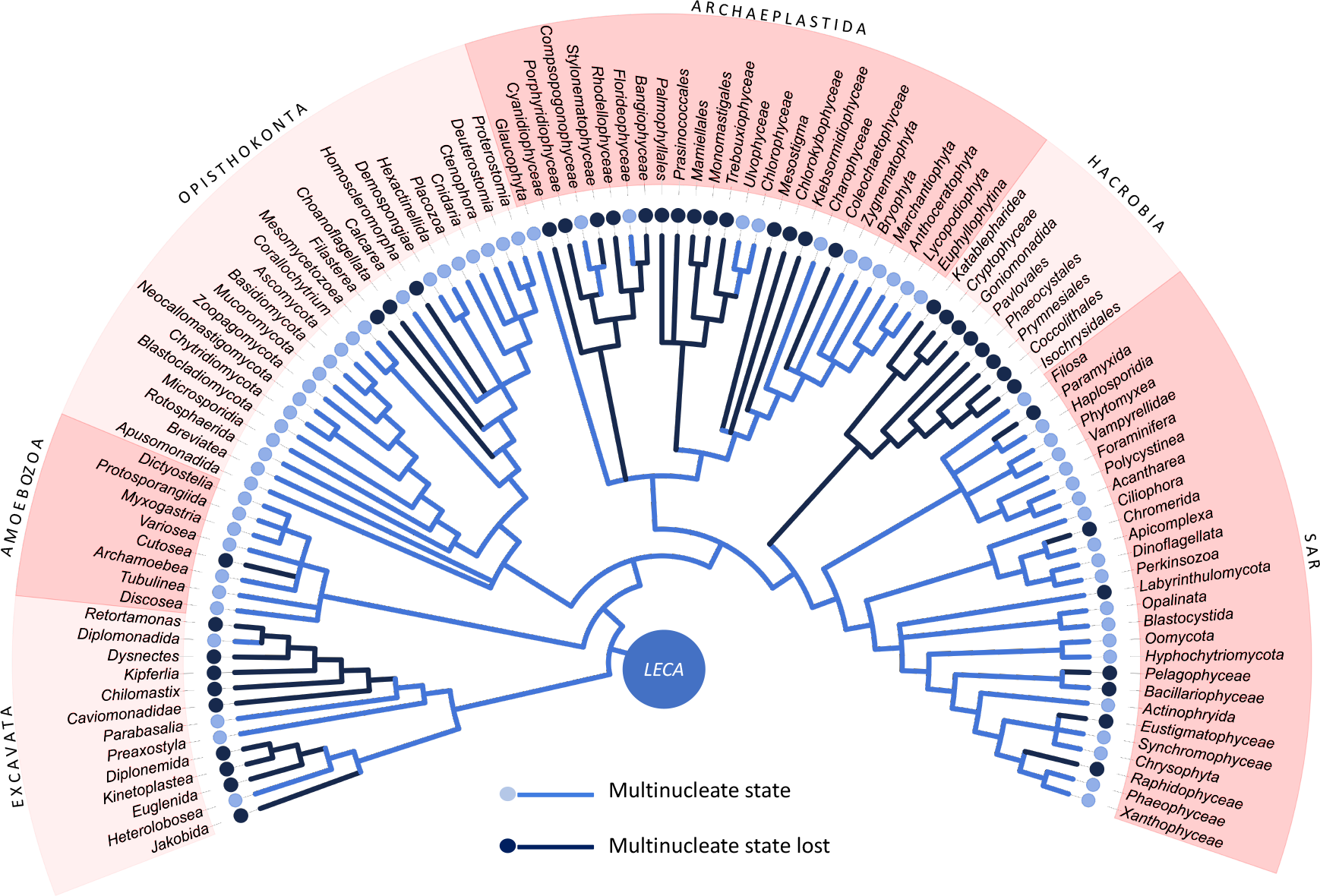
Ancestral state reconstruction for nuclear organization in eukaryotes. Presence and absence of the multinucleate state in members of the respective group is indicated. Resolution of the branches polytomy *versus* dichotomy) does not alter the outcome of the ancestral state reconstruction, nor does position of the root on the branches leading to Amoebozoa, Excavata, or Opisthokonta. LECA was a multinucleate, syncytial cell, not uninucleate (see **Supplemental Figure 2**). Together with mitochondrion^23,54^ and sex^3,31^, the multinucleate state is ancestral to eukaryotes and fostered accumulation of duplications (see text).

The syncytial state allows the independent evolution of nuclei as units of selection^31^. Yet intrasyncytial complementation of defective nuclei only operates if defective nuclei are physically mixed so that the products of mRNA from different nuclei can interact. Mixing of nuclei is characteristic of eukaryotes with syncytial hypha^32^, it requires motor proteins that pull organelles along cytoskeletal elements. Motor proteins are a eukaryote-specific invention^33^ and are noteworthy among LECA’s duplicates in two respects. First, the protein with the most duplications found in LECA is a light chain dynein with 12 duplications (**Supplemental Table 3**), in agreement with previous studies of dynein evolution that document massive dynein gene duplications early in eukaryote evolution^34^. Second, 10 of the 20 genes encoding cytoskeletal functions that were duplicated in LECA (**Supplemental Table 3 and 4**) encode dynein or kinesin motor proteins. In contrast to genes transferred from the mitochondrion, where the transfer mechanism itself promotes duplications^15–16^ (**Fig. 3**) in a selectively neutral manner, duplicates of archaeal-derived or eukaryote-specific genes would require selection to be fixed as diversified families. The selective pressure fixing duplications of motor proteins is evident: nuclei encoding motor proteins would mix more effectively than those lacking motor proteins, leading to greater physical intrasyncytial dispersal of nuclei expressing mRNA for motor proteins and increased fitness of nuclei encoding them. Individual nuclei in a syncytium have properties of individuals in population genetics^31,32^, serving as units of selection both for the origin of cytonuclear interactions, and at the level of uninucleate, mitochondriate spores for the generation of mitotic progeny as the first typically protist-like cells. The syncytial state presents a viable intermediate state in the transition from prokaryote to eukaryote genetics. Gene duplications in LECA uncover an early origin of mitochondria and record the onset of the eukaryotic gene duplication process, a hallmark of genome evolution in mitosing cells^17–21^.

## Methods

### Dataset preparation

Protein sequences for 150 eukaryotic genomes were downloaded from NCBI, Ensembl Protists and JGI (see **Supplemental Table 5** for detailed species composition). To construct gene families, we performed an all-vs-all BLAST^35^ of the eukaryotic proteins and selected the reciprocal best BLAST hits with e-value ≤ 10^−10^. The protein pairs were aligned with the Needleman-Wunsch algorithm^36^ and the pairs with global identity values < 25% were discarded. The retained global identity pairs were used to construct gene families with the Markov Chain algorithm^37^ (version 12-068). Because in this study we were interested in gene duplications, we considered only the gene families with multiple gene copies in at least two eukaryotic genomes. Our criteria retained a total of 24,571 multi-copy gene families.

### Sequence alignment and gene tree reconstruction

Protein-sequence alignments of the individual multi-copy gene families were generated using MAFFT^38^, with the iterative refinement method that incorporates local pairwise alignment information (L-INS-i, version 7.130). The alignments were used to reconstruct maximum likelihood trees with IQ-tree^39^, using default settings (version 1.6.5), and the trees were rooted with the Minimal Ancestor Deviation method^40^.

### Inference of gene duplication

Duplications in the rooted topologies were identified from all pairwise comparisons of multi-copy genes sampled from the same genome. Given a rooted gene tree with *n* leaves, let *S* the set of species labels for the leaves. For the particular case of multi-copy gene trees there is at least one leaf pair, *a* and *b*, such that *s*_*a*_ = *s*_*b*_. Because the tree is rooted it is possible to identify the internal node corresponding to the last common ancestor of the pair *a* and *b*, where the internal node corresponds to a gene duplication. For each gene tree, we performed pairwise comparisons of all leaf pairs with identical species labels to identify all the internal nodes corresponding to gene duplications. This approach considers the possibility of multiple gene duplications per gene tree and minimizes the total number of gene losses. Genes descending from the same duplication node form a paralogous clade (**Supplemental Figure 1**). It is possible that not all the species in a paralogous clade harbor multiple copies of paralogs, due to gene loss. Therefore, variable copy-number of paralogs among the species present in the same paralogous clade is indication of, at least, one gene loss event. We summarized the duplication inferences from all the trees by evaluating the distribution of paralogs descending from duplications across the six eukaryotic supergroups (**Fig. 2**).

### Identification of homologs in prokaryotic genomes

For identification of homologs in prokaryotes, we used protein sequences from 5,524 prokaryotic genomes (downloaded from RefSeq^41^, see **Supplemental Table 6**) and compared those against the eukaryotic genes using Diamond^42^ to perform sequence searches with default parameters. A eukaryotic gene family was considered to have homologs in prokaryotes if at least one gene of the eukaryotic family had a significant hit against a prokaryotic gene (e-value < 10^−10^ and local identity ≥ 25 %).

### Ancestral reconstruction of eukaryotic nuclear organization

Ancestral state reconstructions were performed on the basis of a morphological character matrix, using maximum parsimony as implemented in Mesquite 3.6 (https://www.mesquiteproject.org/). The reference eukaryotic phylogeny includes 106 taxa (ranging from genus to phylum level) to reflect the relations within the eukaryotes and reduce taxonomic redundancy. The phylogeny includes members of six supergroups: Amoebozoa (Mycetozoa), Archaeplastida, Excavata, Hacrobia, Opisthokonta, and SAR, and was constructed by combining branches from previous studies^22,43–59^. The nuclear organization for each taxon was coded as 0 for non-multinucleate, 1 for multinucleate or 0/1 if ambiguous according to the literature^22,43,55,59,60–68^ (**Supplemental Table 7**). In order to account for uncertainties of lineage relations among eukaryotes, we used a set of phylogenies with alternative root positions^23,69–71^ (altogether a total of 15 different roots) as well as the consideration of polytomies for debated branches (**Supplemental data**).

## Acknowledgements

We thank the European Research Council (grant 666053) and the Volkswagen Foundation (grant 93 046) for financial support. We thank Nils Kapust, Michael Knopp, Damjan Franjević (Departmnent of Biology, University of Zagreb, Croatia) for helpful discussions.

